# Dynamo: An open-source Python application for quantifying neuronal structural plasticity over time

**DOI:** 10.64898/2026.09.09.750485

**Authors:** Peter William Hogg, Patrick Coleman, Jason Fung, Kaspar Podgorski, Tristan Dellazizzo Toth, Kurt Haas

## Abstract

Neurons exhibit tremendous structural plasticity during the growth of dendritic and axonal arbors in early brain circuit formation, followed by experience-driven structural plasticity throughout life. Advances in labeling and *in vivo* time-lapse imaging allow capture of dynamic structural changes within intact and awake animals, and fluorescent biosensors of neural activity offer opportunities to link structural and functional plasticity. However, the resulting large multi-dimensional data sets are challenging to quantify due to time-consuming tracking of minute structural changes throughout complex neuronal morphologies across time. Here, we present *Dynamo:* an open-source Python application that enables *dynamic morphometries*, the quantitative analysis of morphological changes over time, by streamlining arbor reconstruction, registration of structures across time, and quantitative analyses of growth behavior. Dynamo yields rich characterization of neural structural changes necessary for determining how rapid growth events culminate into long-term patterning, linking structural and functional plasticity, and for identifying underlying molecular mechanisms.

**HIGHLIGHTS:**

- Dynamo tracks individual neuronal structures across 4D time-lapse imaging data
- A Tree-Branch-Point hierarchy preserves element identity as arbors remodel
- Machine learning and manual tools enable fast, accurate 3D/4D reconstruction
- Dynamo links structural dynamics to molecular and neural activity signals

**eTOC BLURB:** Hogg, Coleman, et al. present Dynamo, an open-source Python platform that represents neuronal arbors as identity-preserving Tree-Branch-Point objects across time-lapse imaging series. Combining machine learning-based tracing with manual reconstruction and registration tools, Dynamo enables direct, programmable quantification of dendritic and spine structural plasticity linked to molecular and activity signals.

## INTRODUCTION

The morphologies of brain neurons are intrinsically linked to their computational abilities to receive, integrate, transform, encode, and transmit information. Dendritic and axonal arbors are complex, increasing surface area to accommodate hundreds to thousands of synapses and allowing connections with distant sites. The study and quantification of these morphologies, or morphometries, has been a valuable tool for neuroscientists since Cajal conducted *camera lucida* drawings to classify neurons into subtypes and infer their contributions to circuit functions^1^. However, neuronal structures are not static, but rather exhibit exuberant growth during developmental stages of brain circuit formation^23^, and dynamic structural rearrangements associated with learning and memory in the mature brain^4^ ^6^. Quantification of these structural changes, or dynamic morphometries, is necessary for understanding how individual neurons achieve their mature morphologies and patterns of synaptic connectivity to form functional networks^5^. These measures are critical for linking molecular mechanisms to structural changes and identifying developmental constraints on circuit structure and function^7^. In mature circuits, dynamic morphometries discerns the potential, limitations, and mechanisms of plasticity.

While advances in sparse neuronal labeling and *in vivo* time-lapse imaging now allow routine acquisition of 4D neuronal growth image data sets^6,8,9^, dynamic morphometries analyses have been hampered by the difficulty of tracking minute changes throughout complex neural structures over time. Extensive efforts have been directed towards development of computer-assisted digital reconstruction and quantification of single time point 3D image stacks for general neuronal morphometries, such as the TREES toolbox^10,12^.

Since manual drawing can be exceptionally time-consuming, automated software approaches have made important contributions by significantly increasing the speed of generating reconstructions, including the BigNeuron project^13^, Vaa3D^14^, and others including commercial software packages^15^. Following digital reconstruction, morphometries can be automatically^15^ calculated^16,18^, and open-source data structures allow archiving and sharing of digitized morphologies^19^. However, current approaches designed for static, single time point images are insufficient for tracking neuronal growth, since they are unable to identify and track changes of individual structural elements across time for sophisticated dynamic morphometries analytics.

To address this, we developed Dynamo, an open-source platform based on representing a neuronal arbor as a persistent hierarchical object where individual structural elements keep their identity across the time series. Dynamo enables reconstruction with automated or manual tracing features, or from an imported SWC file, registering and tracking the arbor so that growth, plasticity, and function can be quantified over time. We demonstrate the application of Dynamo for rapid machine learning-based automated reconstruction of several neuron types, including tracking and quantification of rapid structural changes during dendritic arbor growth in *Xenopus* tadpoles, and tracking spine dynamics of pyramidal neurons in the primary visual cortex of mice. Further, we demonstrate the utility of Dynamo for investigating underlying molecular mechanisms by tracking subcellular markers along with growth, and for linking structure to function by pairing spine morphological plasticity with neural activity recorded using calcium and glutamate biosensors.

## RESULTS

Dynamic morphometries requires a time series of 3D image stacks of single- or sparsely labeled neurons in order to correctly trace dendritic or axonal processes (Fig. 1). To demonstrate Dynamo’s functionality in streamlining dynamic morphometries for studies of dendritogenesis, individual newly-differentiated brain neurons in the developing albino *Xenopus laevis* optic tectum were fluorescently labeled using single-cell electroporation^8,20^ for delivery of plasmid DNA expressing the genetically-encoded fluorophores EGFP or LifeActEGFP. Two days following transfection, awake tadpoles were immobilized and 3D volumes encompassing the soma and full dendritic arbors of the labeled neurons were imaged by *in vivo* two-photon microscopy using serial optical sectioning with a 1.5 *pm* step size. A time series of full-volume image stacks was repeatedly captured at 5-min intervals over 120 min for EGFP, and 3-min intervals for 30 min for LifeActEGFP. Previous studies of *Xenopus* tectal neuron dendritogenesis find that 5-min interval imaging is sufficient for recording all process additions, eliminations, and accurate measures of growth motility (rates of process elongation and reduction)^2^. Longer imaging intervals from hours to days is required to capture large-scale patterning to form the mature dendritic arbor^2^. To demonstrate Dynamo’s ability to track these larger changes, additional tectal neurons labeled with EGFP were imaged daily for three days as their dendritic arbors elaborated.

**Figure 1.**
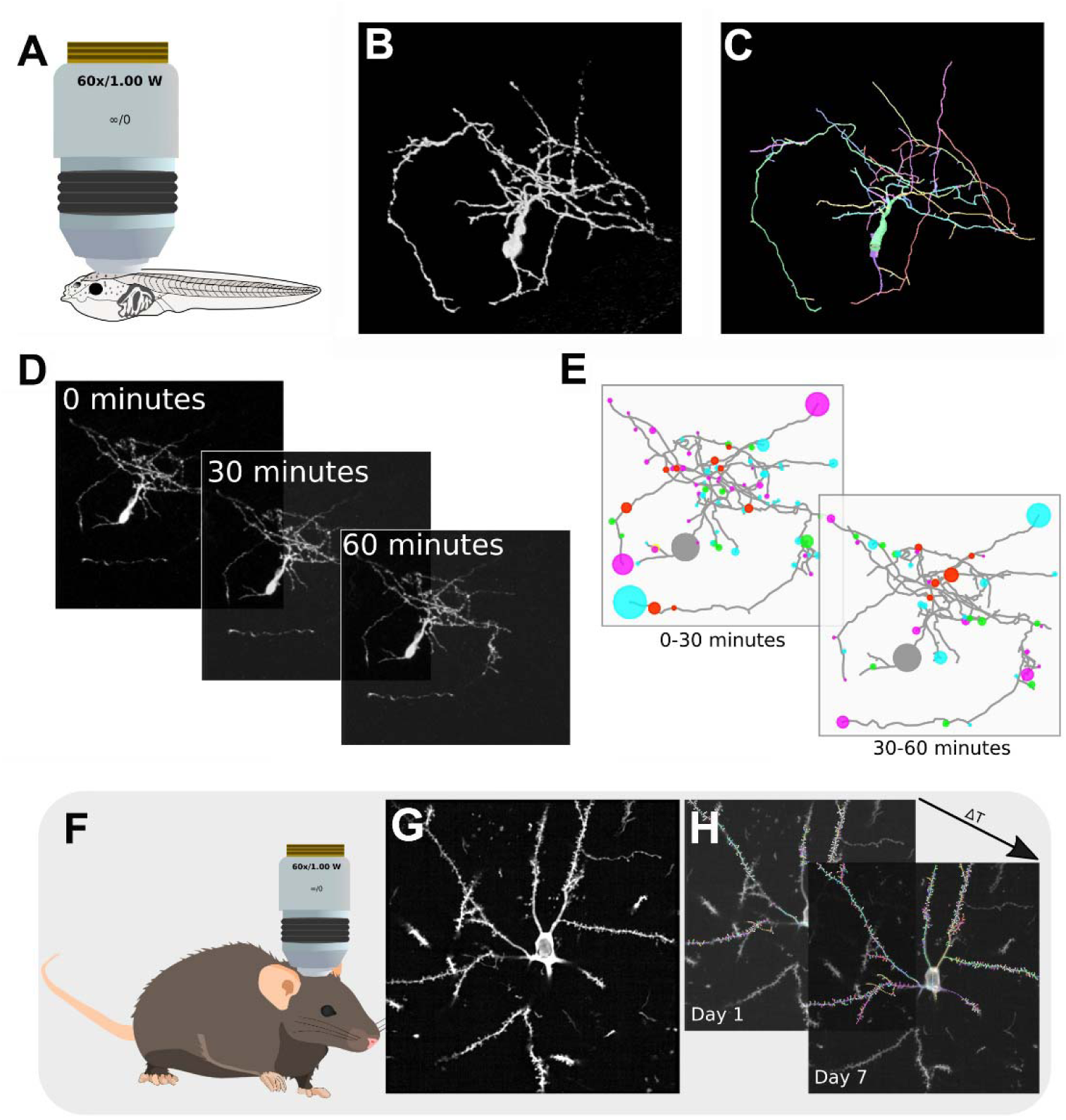
Dynamic morphometric analysis workflow in Dynamo. **A**, *In vivo* two-photon time-lapse imaging of a 3D volume encompassing one neuron in the transparent albino *Xenopus* optic tectum labeled by single-cell electroporation for expression of EGFP (Video S4). B, 2D projection of 3D volume encompassing the entire dendritic arbor, soma, and proximal axon. C, Dynamo automated reconstruction of neuron in b, with color denoting separate Branch Objects. **D,** Repeated imaging of 3D volume at 30-min intervals to capture growth. E, Computer-assisted reconstruction and spatial registration of neural structures across time points, with colored circles at the tips of protrusions displaying new additions (green), eliminations (red), elongations (cyan) and reductions (magenta), and circle size depicting change in process length. See Sup. Fig. 4 for dendrograms. **F,** *In vivo* two-photon time-lapse imaging of pyramidal neuron dendritic spine plasticity in mature mouse visual cortex, with sparse fluorophore expression. G, 2D projection of a volume of visual cortex encompassing the soma and proximal dendrites of a pyramidal neuron. H, Repeated imaging of the same volume at 7-day interval for tracking spine morphology changes over time. Branch Objects including dendritic branches and spines are assigned random colors representing unique identifications that are consistent across time points.

To visualize the dynamic morphometries of dendritic spine plasticity in the mature brain, pyramidal neurons in mouse primary visual cortex were co-labeled with the genetically-encoded fluorescent biosensors iGluSnFR-SF-VenusA184S to detect extracellular glutamate, and jRGECO1a for intracellular calcium, allowing dual recording of visually-evoked pre- and post-synaptic responses, respectively. *In vivo* two-photon volume imaging of the basal fluorescence of the biosensors throughout the pyramidal neuron dendritic arbor allowed capture of the structure of large numbers of spines. Volumes were imaged twice with a 7-day interval to capture neuronal structural changes, along with rapid imaging of visual-evoked pre- and post-synaptic activity to correlate structural plasticity with synaptic function^21^.

### Importing 3D Image Volumes into Dynamo

A time series of 3D image stacks from time-lapse volumetric imaging are imported into a Dynamo project sequentially as an ordered collection of multi-color channel images (Fig. 1; Video S1). File formats currently supported are tiff (.tif), MATLAB (.mat) and Zeiss’ LSM (.czi). Each time point in the project is represented as a volume of fluorescence intensity data in a 4D array, with data in the order of: color channel, Z stack, Y pixel, X pixel. Each of the loaded single time point volumes within a time series are displayed in separate windows, allowing for simultaneous viewing of morphological changes across the time series that can be viewed slice-by-slice, rendered in 3D, or as a 2D maximum projection. Dynamo also allows users to quickly alternate between grayscale and perceptually uniform colormaps to better visualize changes in imaged signal intensity.

### Machine Learning-Based Automated 3D Neuronal Reconstructions

Dynamo supports both semi- and fully-automated neuronal reconstructions using a machine learning-based image segmentation approach. A Deep Residual U-net, trained on manually curated reconstructions, classifies each voxel of the reference image stack as neural or non-neural (Fig. 2). The trained U-Net can automatically identify fluorescent labeled neurons as foreground from the background, and further classify structures as soma, dendrites, and all other biological signals. Using the center of mass of the soma pixels, a tree can be built using the remaining classified pixels. The segmented 3D neuronal structure of the dendrites is skeletonized to generate points for the Dynamo tree that include a unique identifier, 3D position, and connections to other branches, all ordered as a tree rooted in the center of the soma. Examples of semi-automated tracing from a user-provided root node are shown in screenshots from the Dynamo UI (Fig. 2G). This approach was evaluated against expert-reviewed ground-truth reconstructions derived from the same imaging system used to generate the training data, as well as against external datasets. The computer vision-based reconstruction method demonstrated performance comparable to other state-of-the-art methods, such as the Vaa3D APP2 tracing algorithm (Sup. Fig. 1)^14^. However, Dynamo and Vaa3D differ in performance depending on the input image, and Dynamo consistently overestimates, and Vaa3D underestimates the number of branch tips (Sup. Fig. 1).

**Figure 2.**
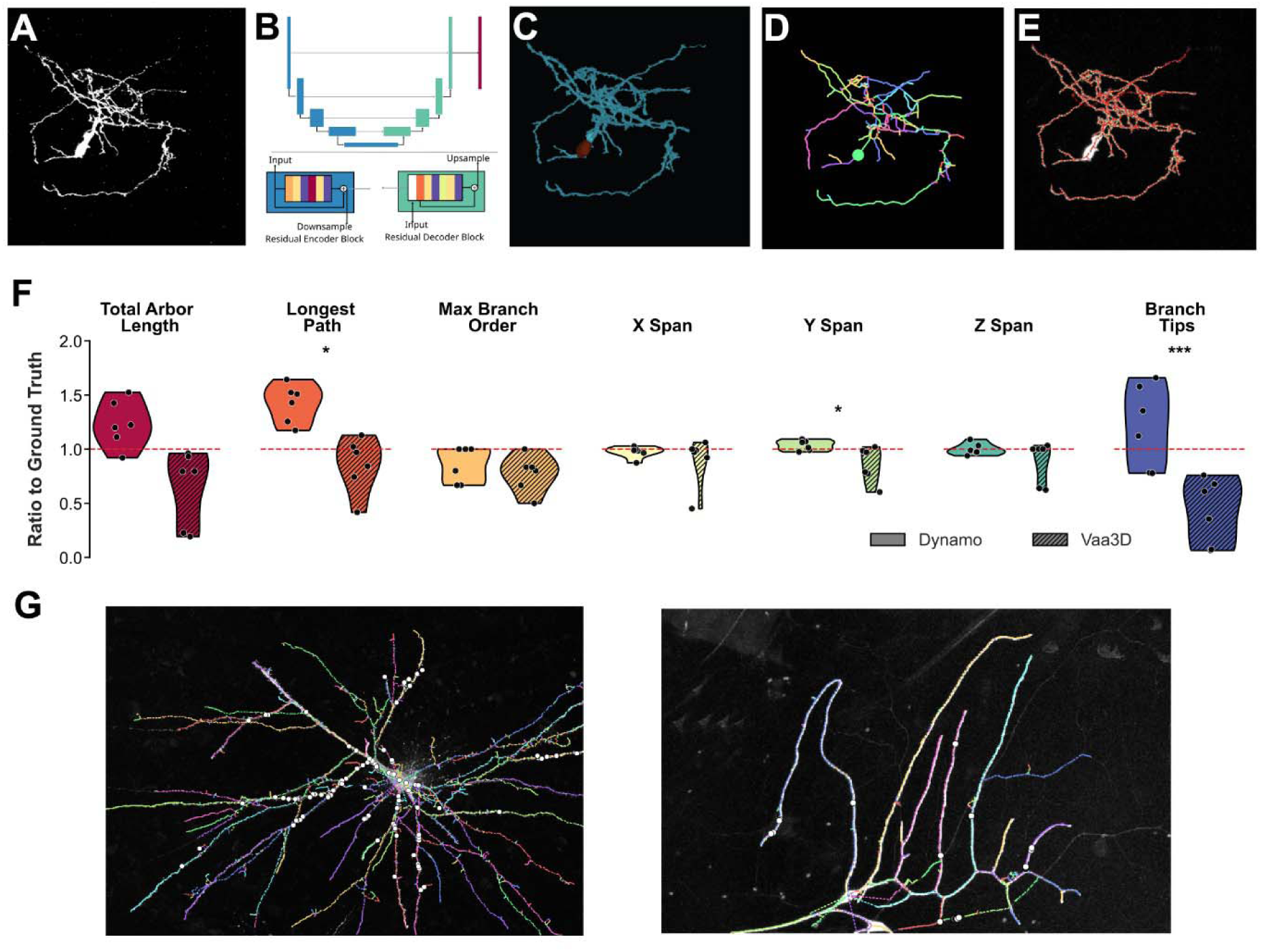
Machine learning-based neuronal tracing using Dynamo. **A,** Reference image of a tectal neuron. B, Schematic representation of a 2D U-Net architecture. C, Pixel classification highlighting the soma *(brown)* and dendrites *(blue).* **D,** Reconstructed neuronal tree generated by Dynamo. E, Overlay of ground truth reconstruction and machine learning-based trace. F, Quantitative comparison of traces produced by Dynamo and Vaa3D, normalized to the ground truth tectal neurons (N = 5 neurons). G, Example overlays of Dynamo reconstructions using semi-automated tracing of a human and a Drosophila neuron (left to right), images retrieved from^43^. Statistical tests: paired t-test or Wilcoxon signed-rank test. *p < 0.05, **p < 0.01, ***p < 0.001.

While this machine learning-based approach offers significant advantages, it has limitations, as tracing accuracy is highly dependent on the similarity of the input images to the training set. To enhance Dynamo’s flexibility, new pixel classifiers can be trained and integrated. For example, widefield image stacks of biocytin-filled neurons can be reconstructed using a U-Net trained to classify neuronal and background pixels (Sup. Fig. 2). As this U-Net cannot identify the soma it requires users to provide the root node’s placement before tracing begins.

In addition to automated 3D neuronal reconstruction, Dynamo also incorporates two manual tracing modes for digital reconstruction, *Skeleton* and *Radii,* with digitized neurons linked to each 3D reference image stack (Fig. 3; Video S2). In the *Skeleton* mode, neuronal structures are reduced to a wire frame, with points placed in the center of branches. Alternatively, the *Radii* mode preserves width at each point (for processes, soma, spines, etc.), which is automatically determined using the intensity values from the reference image volume.

**Figure 3.**
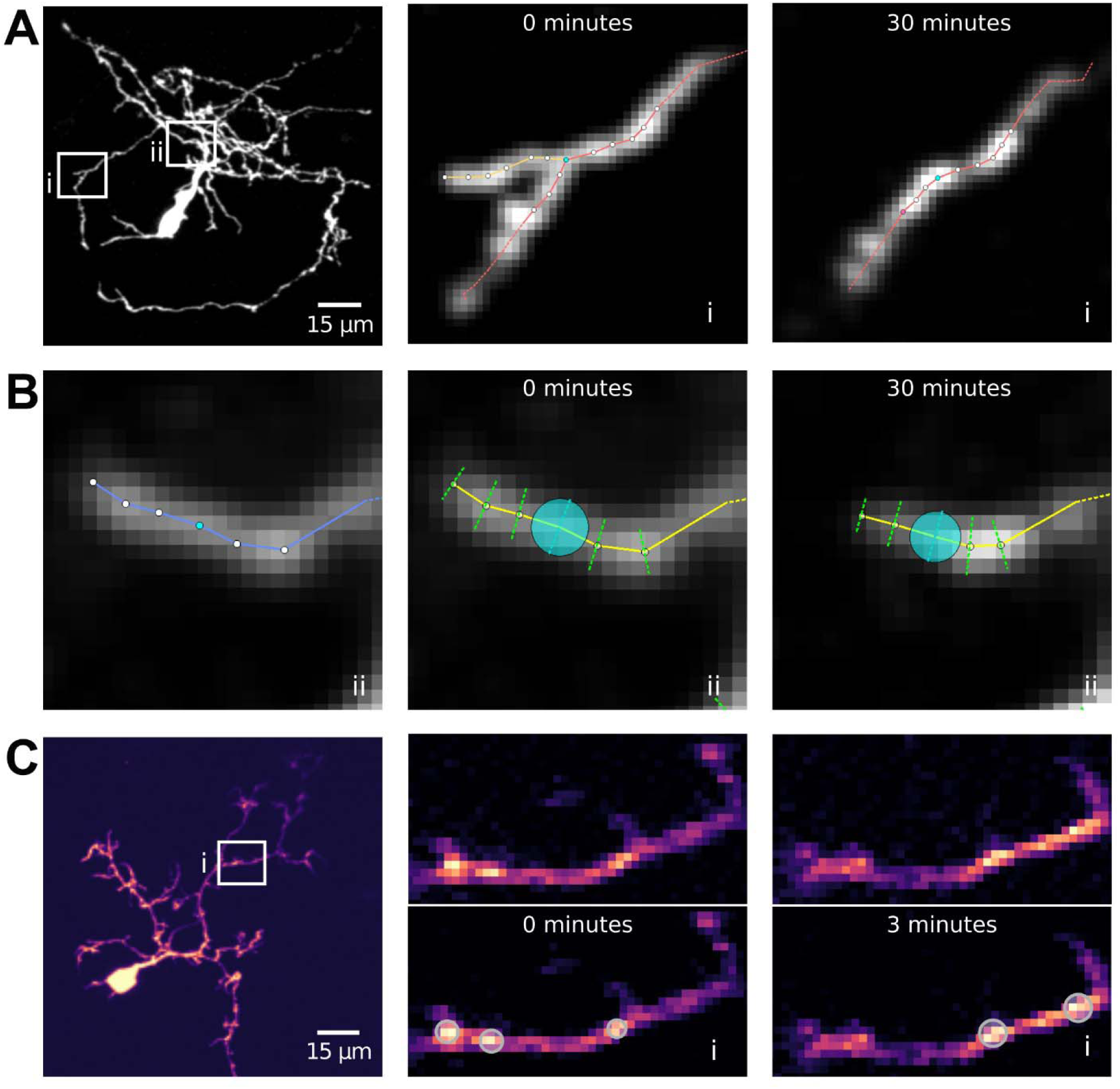
Neuronal drawing and editing using Dynamo. Drawing modes include: **A,** *Skeleton* tracing mode for digitizing arbor structures by reducing branches to a series of points at the midpoint of all processes. **B,** *Radii* tracing mode for digitizing neuronal structures while preserving processes diameter information. **C,** *Puncta* mode allows tracking the position and size of objects stored separate from, but linked to the digitized tree hierarchy. Here, puncta of fluorescently tagged-actin LifeAct-EGFP are identified and tracked over time.

### Reconfiguring 3D reconstructions into identified objects

For both the *Skeleton* and *Radii* modes, digitized neuronal structures are organized into a hierarchy of Python objects, with methods allowing for easy spatial analytics and referencing. The top of the hierarchy is the Tree Object, which is the parent of a collection of Branch Objects composed of an ordered series of Point Objects. Each object has a unique identification value linking its spatial location in the volume and its relationship to other objects in the hierarchy. In the *Radii* mode the diameters of all the neuronal processes are visualized and editable, and these values are stored in the Point Object similar to the SWC file format. The hierarchy of objects starts at one Point Object, the root node, which is generally placed in the center of the cell body or the start of a dendritic or axonal branch depending on the scope of the reconstruction.

Dynamo can also import neuronal reconstruction traces stored in SWC files that have been generated by other drawing tools such as Vaa3D^14^ and Mozak^22^. The structural information in the SWC file format is used to populate the Points, Branch and Tree Objects in the Dynamo hierarchical data structure (Sup. Fig. 3; Video S3). Since the format of SWC files often results in an undesirable branch order upon initial import, Dynamo can solve this by reparenting the branches by path length or angle automatically resulting in structures optimized for calculating morphometries such as the longest path length or branch order.

### Digital Neuronal Reconstruction of 4D Time Series

Once the reconstruction of the first time point of a 4D time series has been completed (Video S4), additional image volumes for subsequent time points can be added to the Dynamo project file. The digitized neuronal structure from the initial time point is imported to, and used as a base template for faster digitization of the next time point. This results in a new Tree Object that has been populated by the values from the importation source, and can be adjusted to accommodate growth changes, including extensions, retractions, or eliminations of existing processes, or new additions. This process of loading additional volumes, importing the previous time point neuronal trace, and updating the reconstruction for the new time point is repeated until the entire time series is reproduced digitally.

### Registration of Arbor Element Identities Across a Time Series

Sophisticated metrics of growth are dependent on tracking the identity of neuronal components across a 4D image stack time series^5^. To identify sites of structural change and to ensure fidelity of object identification, Dynamo implements three structural registration methods that align digitized arbors between time points, and synchronizes the unique IDs for each object in the hierarchy: *Image, ID* and *Manual Registration*.

*Image Registration* should be utilized after importing a Tree Object from the preceding time point to a subsequent image data set. Dynamo algorithms can adjust the point positions in the imported reconstruction to accommodate for image drift and morphological changes due to growth and plasticity. *Image Registration* is a novel image-based algorithm that utilizes both Point Object positions and voxel intensities between two successive image stacks (Fig. 4 A-C). When conducting digitization in Dynamo and a reconstruction is transferred to a subsequent image stack, Dynamo conducts an automated adjustment of the Point Object positions to best match the voxel intensities in the new image stack to accommodate for image drift. Pairs of points are examined recursively by comparing a local sub-volume (21 x 21 x 9)px centered around their voxel positions. These images are extracted, then flattened to 2D maximum intensity Z projection, and the optimal affine transformation (translation and rotation) is found to align the sub-volumes. If a close match is found, the (X, Y) translation is applied, and an optimal Z offset is calculated, updating the spatial location of the Point Object in the second tree (Fig. 4 B,C). If no close match is found, the process is repeated with larger (51 x 51 x 9)px neighborhoods, to allow for even greater object motion. If this expanded matching attempt also fails, the points are skipped, and registration continues along the arbor. Once complete, successful matches will have their IDs synchronized, and aligned to locations that most closely match the earlier time point. Unregistered points are highlighted in a different color, to alert the user for manual inspection. This registration process is sensitive to noise levels, arbor complexity, and the amount of drift and rotation of the neuronal structure between consecutive imaging volumes.

**Figure 4.**
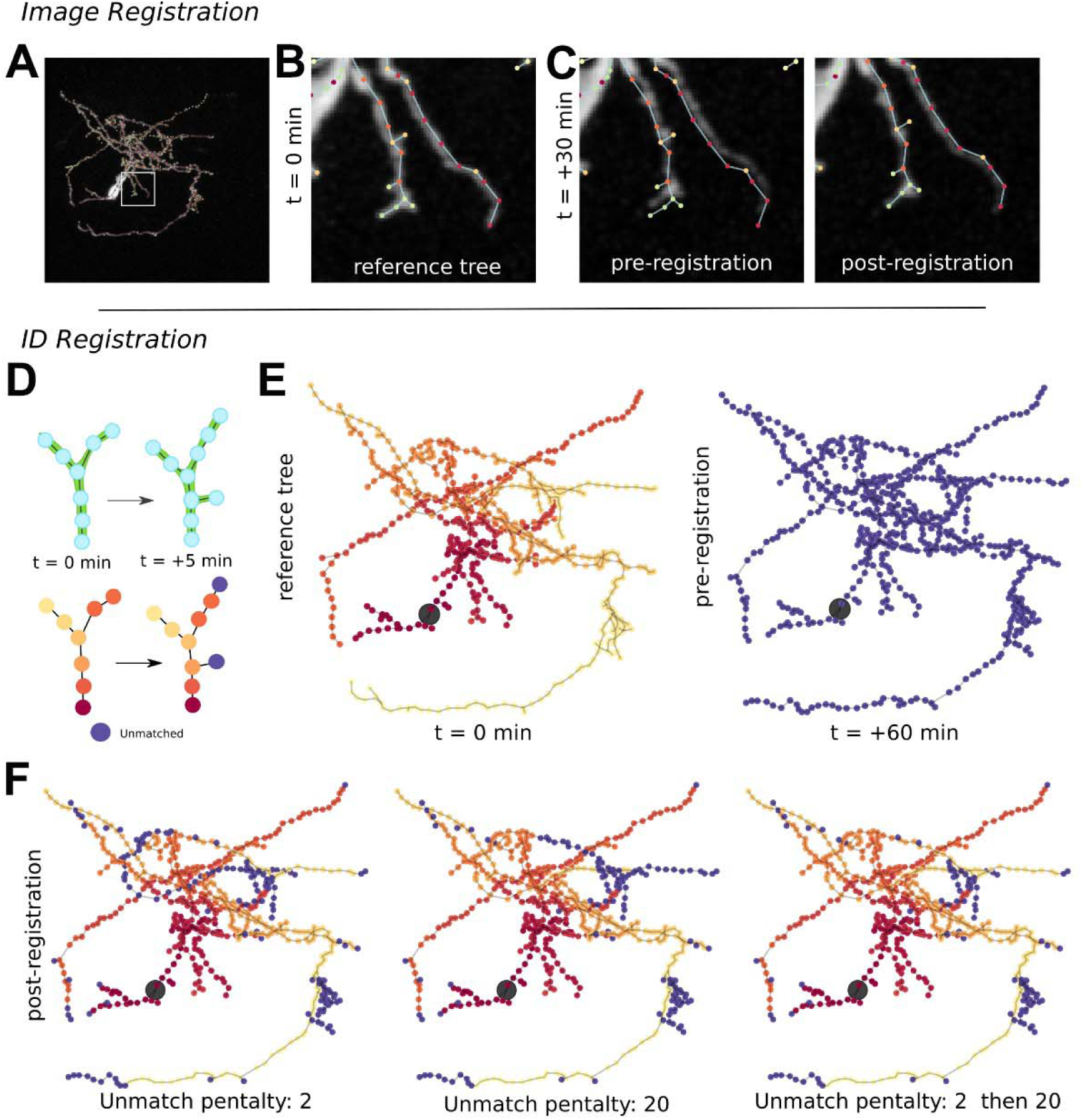
Automated registration methods of digitized neuronal structures across time. Dynamo synchronizes the identification of points across successive time points using 3 registration methods: *Image, ID* and *Manual.* In *Image Registration,* an initial time point neuronal reconstruction **A-B,** is transferred to the subsequent image volume **C.** An automated routine using intensity values in the volume surrounding Point Objects adjusts the image volume in 3D to optimize digital reconstruction-image overlap. **D,** *ID Registration* uses an automatic cost-based algorithm to synchronize Branch and Point Objects identifications across successive time-points. E, Two digital reconstructions from the same neuron imaged at a 60-min interval imported into Dynamo, with each Point Object in the t = 0 reference tree uniquely colored on a colormap, and as yet unmatched Point Object in the t = +60 min tree colored blue. F, *ID Registration* results using an unmatched penalty of 2, 20, and two rounds of registration using both values. The remaining unmatched Point Objects are then reviewed for *Manual Registration* to correct errors, or accept new points due to growth.

A second automated method, *ID Registration,* uses an optimization framework to ensure that IDs of objects are correctly synchronized over time points without altering the position of Point Objects (Fig. 4D-E). This algorithm is particularly important for comparing two Tree Objects that have many Point Objects with unmatched IDs, such as when importing external digital reconstructions from SWC files, rather than conducting reconstructions in Dynamo. To synchronize object IDs between two reconstructions, Dynamo uses the spatial positions of Point Objects and their relationships within the Tree Object hierarchy to find an alignment solution between two sequential Tree Objects that minimizes a cost function^23^. This cost-minimization approach is similar to the Dynamic Time Warping method used to register axonal branches^24^, where the registration cost is calculated as the sum of the real-world distance between registered pairs of Point Objects, as well as a scaled cost for all unregistered Point Objects to account for dynamic growth and deletions (Sup. Equation 1). The scaled cost is defined by the difference of each Point Object’s position relative to their paired predecessor, to be resilient to any translation of the neuron between time periods. By combining the two penalties (distance, and unmatched counts) with a scaling factor, the user can balance matching Point Objects and distance to control how far a point can have moved in 3D space before it should be considered ‘new’. Using a recursive algorithm to find the optimal alignment between trees, the efficiency of finding the lowest-cost alignment can be calculated as O(N^2^B!), where N is the number of points per tree, and B is the average number of branches off each point^25^. Dynamo uses a default scale factor of 10 *gm,* i.e. a point can move 10 *gm* before it will be considered ‘new’, however this threshold can be altered. Once complete, the user is presented with the pairing that produces the minimal cost, and the option to accept or reject that registration.

If these automated methods of registration fail, Dynamo supports a third, *Manual Registration* approach, in which user-selected points across multiple arbors can have their unique object IDs synchronized.

### Error Correction

Errors are common in digital reconstructions due to multiple factors, including the complexity of neuronal structures, small protrusions near the optical resolution limit, the use of low excitation power required to limit phototoxicity and bleaching, as well as inherent limitations of automated methods. Identifying mistakes in reconstructions is particularly critical in dynamic morphometries, since an error early in a time series may persist throughout subsequent time points, compounding to generate large discrepancies in analyses. Therefore, manual correction is typically required, and is a rate-limiting step in neuronal reconstruction pipelines^26^. To streamline error-correction, Dynamo has a comprehensive set of rapid manual and automated editing tools that synchronize corrections across future time points. Neuronal reconstructions either created in Dynamo or imported are overlaid on top of their reference image volumes and can be interrogated together in 3D using the Python visualization toolkit Napari^27^ (Video S5). Common errors arise from misclassification of points to structures, such as new branches being erroneously assigned as extensions of their parent axon or dendrite, rather than identified as a new protrusion. In addition, small extensions or reductions of processes may not be accurately tracked, or image noise may be incorrectly misassigned as a neuronal structure. Such errors can be readily detected by visual inspection and manually reassigned. Dynamo also applies an automated error detection function that highlights unlikely morphologies for user review, such as acute angles between connecting points, since branches typically exhibit large radii of curvature.

### Dynamic Morphometric Analyses

Once a time series of 3D volumes is fully reconstructed, registered over time, and reviewed for any errors, 4D quantitative analyses of structural changes can be conducted. Dynamo has in-built dynamic morphometric analyses (Sup. Table 1) directed toward dendritogenesis, axonogenesis, and dendritic spine plasticity, as well as an accessible platform for the development and implementation of novel user-designed measures (Video S6). For all sample types, Dynamo’s inherent ordering of reconstructions into rich Tree, Branch, and Point Object hierarchies with shared IDs allows sophisticated 4D analytics using object-orientated Python programming. Further, multiple visualization options in Dynamo assist in exploration of the processed data. For investigations of neuronal growth, a suite of visualization tools and analytic metrics are designed to track structural changes throughout entire dendritic and axonal arbors including graphs showing locations and measures of growth, changes in total branch length and number, and arbor complexity using 3D Sholl analyses^28^ (Fig. 5B).

**Figure 5.**
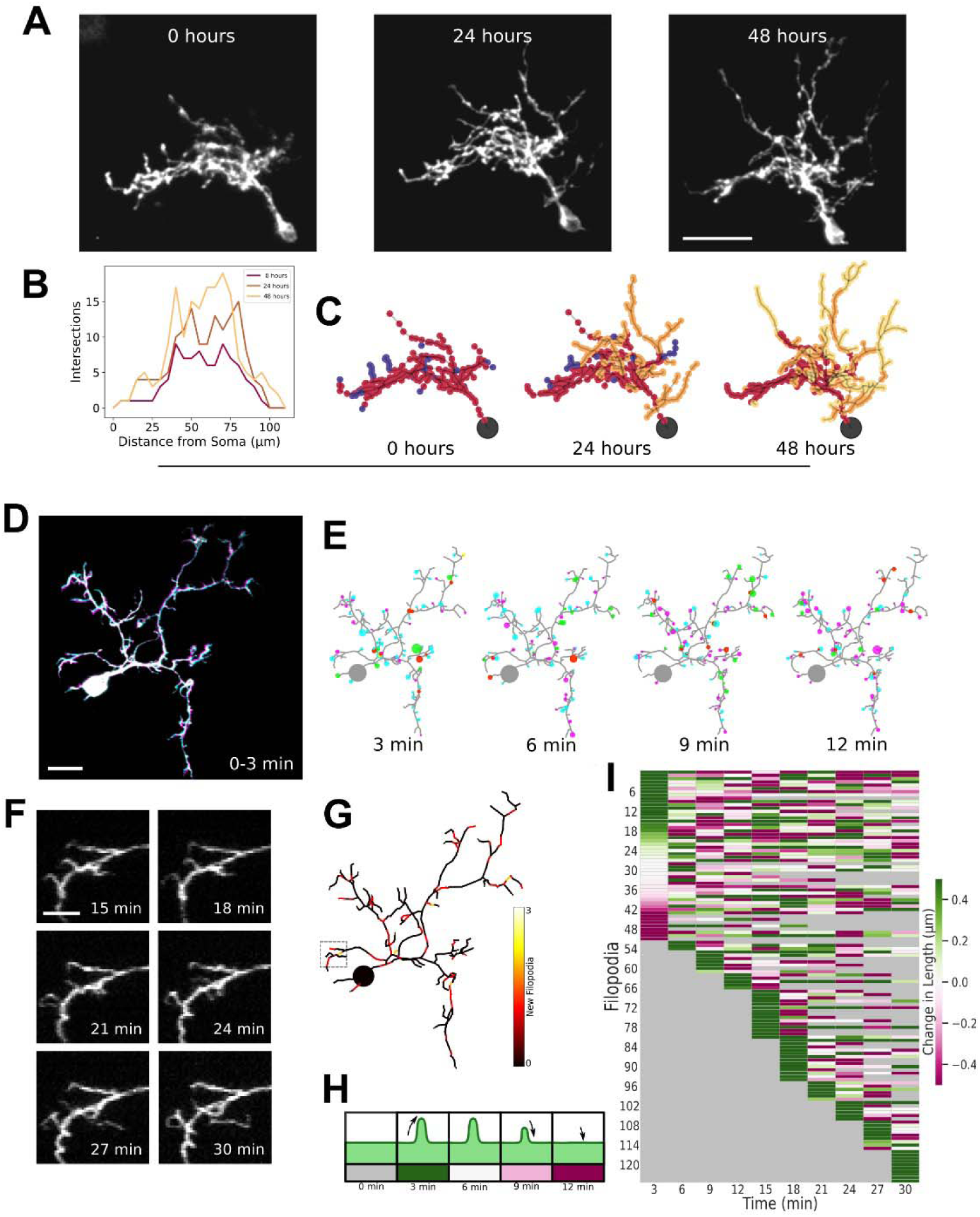
Dynamic Morphometric Analyses. **A,** *AXenopus* tectal neuron imaged over days shows major branch remodeling (scale bar: 25μm). B, Sholl analysis (Dynamo) of neuron in **(A). C,** Dynamo reconstructions color-coded by first appearance (0 h, red; 24 h, orange; 48 h, yellow); pruned branches are blue. D, Short-term remodeling revealed by overlay of two time points (30min interval): t=0 in magenta, t=3 min in cyan, with stable overlap in white (scale bar: 15 μm). **E,** Motility diagram of **(D)** over five time points: soma/stable arbor in gray, protrusion tips indicating additions *(green),* eliminations *(red),* elongations *(cyan),* or reductions (magenta), f, Six Z-projections of a dynamic arbor region over 15 min (scale bar: 5 *gm).* **G,** An arbor heatmap of **(D,F)** shows clustering of addition events. **H-I,** A motility heatmap depicts length changes *(green=elongation, magenta=reduction,* white=stable) and survival of all filopodia (<10 *gm)* every 3 min over 30 min.

Dynamic morphometries can be performed at different temporal and spatial scales which can be accommodated by Dynamo. Larger dendritic arbor remodeling can be observed over several days (Fig. 5A-C), while the changes occurring at small protrusions can be seen on the order of minutes (Fig. 5D-H). Dynamo can handle tracking changes on both of these scales as seen in the example data. With user-defined branch length thresholding, dendritic elements can be separated into distinct groups for analyzing longer branches and shorter filopodia and spines separately. Since small processes are typically the most dynamic structural elements across short time intervals, Dynamo allows branches below a user defined length threshold to be analyzed as a unique group. This allows spine and filopodia growth behaviors, including addition and elimination rates as raw numbers or as density (changes per unit branch length) to be readily tracked, analyzed, and visualized locally, or across the entire neural structure.

For analytics of dendritic spine morphological changes, Dynamo can classify small protrusions as filopodia or spines of different maturational stages based on their length, head, and neck width, all relevant data is stored in the Branch and Point Objects^29^ ^30^ (Fig. 6D). Critically, Dynamo allows tracking of structural changes of the same elements across time, including for metrics of process survival or lifetime, clustering, or other spatial measures linked to previous time point behaviors, such as the location of new process additions in relation to existing structures, and transitions of the classification of neuronal elements, such as the conversion of filopodia into spines (Fig. 6G).

**Figure 6.**
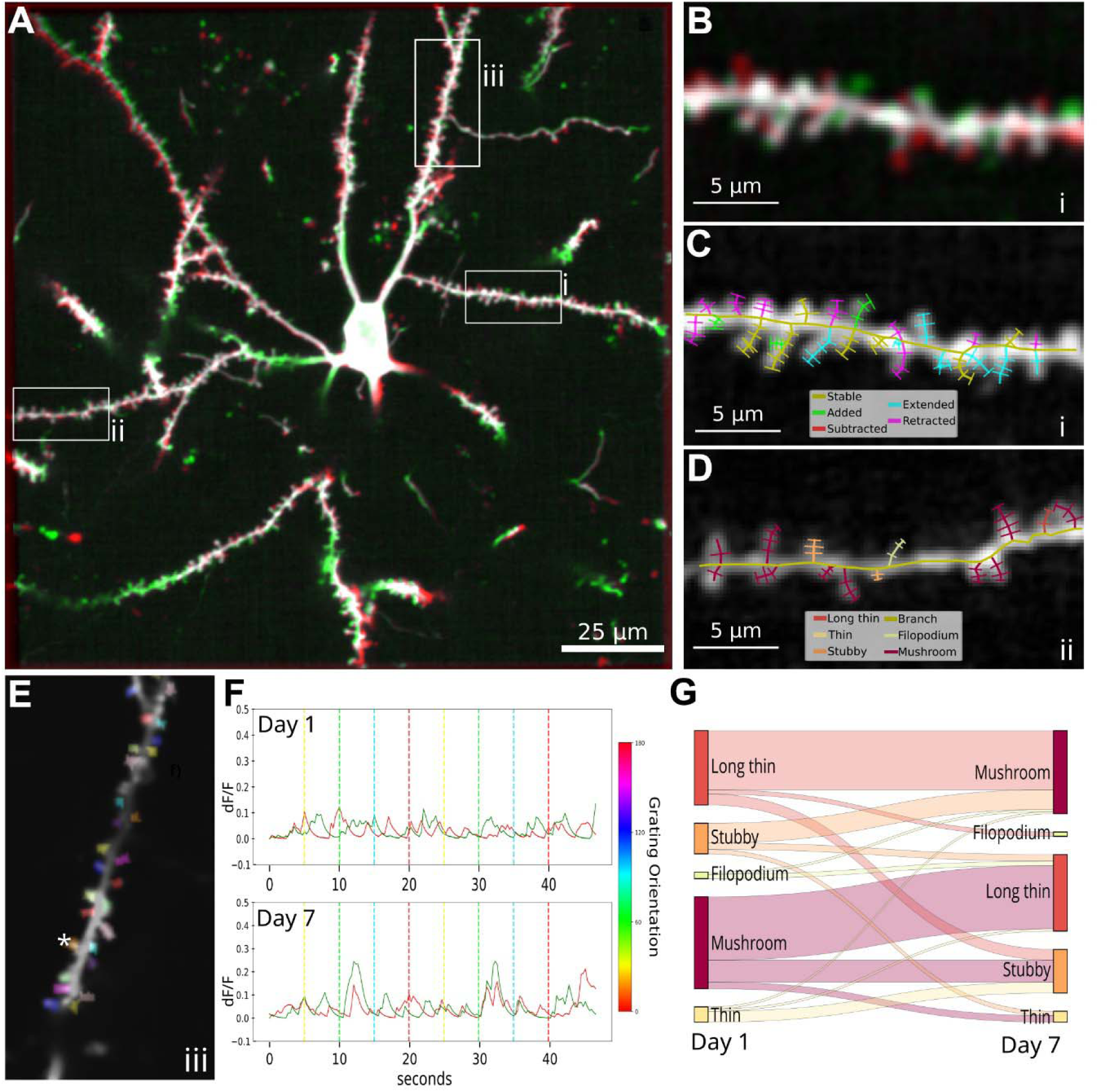
Comprehensive structural and functional analyses of dendritic spines across time. **A,** A pyramidal neuron in the mouse visual cortex imaged *in vivo* at day 1 *(green)* and 7 *(red),* with stable structures in *white.* **B,** Enlargement of region i in **(A). C,** Dynamo reconstruction of dendritic spines in region i, with growth behavior across successive time points indicated by line color. **D,** Dynamo reconstruction of dendritic segment in region ii using radii measures to identify and classify spines, e, Enlargement of region iii with colors depicting regions of spines sampled for fast activity imaging, **f** Activity traces from the dendritic spine marked with * in (E), with calcium signal from jRGECO1a *(red)* and glutamate signal from iGluSnFR-SF-VenusA184S *(green).* Dotted lines indicate presentation of moving-bar visual stimuli, with color indicating bar orientation. **G,** Sankey plot displaying spines transitions between day 1 and 7, excluding spines that did not transition between types.

Basic data and metrics can be exported in csv file format, which can be imported into any statistics software (Sup. Table 1). Given the diversity of experimental questions, model systems and time-varying data sets, Dynamo also includes a rich repertoire of utility functions allowing users to script custom analytic tools for specific experimental needs. As an open-source Python application, users can then share new Dynamo analytics with the larger community.

We demonstrate the ease of designing and implementing new analytics in Dynamo by incorporating metrics for tracking subcellular cytoskeletal markers, and rapid time-lapse imaging of two measures of neural activity, which are linked to structural dynamic morphometric measures. To assist novel feature incorporation, Dynamo has a modality termed ‘Puncta’ (i.e. points in 3D space), where users can identify objects independent of the Tree Object hierarchy and track their movement and size over time. Such markers can be intracellular (e.g. mitochondria, or the synaptic marker PSD-95), extracellular, or on the cell surface (e.g. glutamate receptors). Puncta locations and radii can be drawn manually (Fig. 3), and analysis options are provided to calculate the size, average and total puncta intensity, as well as identification of the closest morphological element, to relate puncta behavior to neural structural changes (Sup. Table 1).

We also show the utility of Dynamo for linking neuronal function to structural growth and plasticity by pairing signals from fluorescent biosensors of activity, including intracellular calcium (jRGECO1a) and extracellular glutamate (iGluSnFR-SF-VenusA184S) with morphological changes over time (Fig. 6; Sup. Video S7). This is supported by importing activity traces using the open Neurodata Without Borders time series data structure^31^. Dynamo can display the raw activity traces and perform several common processing filters, such as ΔF/F0 and non-negative deconvolution^32^. Activity can also be visualized as color-coded heatmaps at each point throughout the reconstructed neuronal structure, or as individual activity traces at all, or a region encompassing multiple sampled points, and across a time series. Dynamo analytic tools can be applied to determine the relationship between local activity and morphological changes over time^33^. For example, the structural dynamics of the dendritic spines of a pyramidal neuron in the mouse visual cortex can be combined with measures of activity across its dendritic arbor (Fig. 6). Dynamo reconstructions can be used to generate and track neural activity in regions of interest at specific spines using the unique IDs of the Branch Objects in the Dynamo project file. This allows the activity of individual dendritic spines to be compared over multiple imaging sessions for linking changes in spine tuning properties to their structural plasticity across time (Fig. 6E-G).

### Dynamo Paired with Biophysical Simulations

As a Python package, Dynamo can be integrated into external analysis pipelines dependent on dynamic neuronal morphologies. Here, we demonstrate how Dynamo can pair *in vivo* functional activity and morphological changes acquired across multiple imaging sessions with an established biophysical model of a visual cortical pyramidal neuron implemented in the neuronal simulation and modeling software NEURON^34^. Calcium imaging data from the pyramidal cell in Figure 6 were used to generate synaptic input patterns for simulations of the neuron’s membrane potential across two imaging sessions. For demonstration purposes, the simulation was restricted to a single reconstructed dendritic branch. The branch and spine morphologies were tracked across imaging sessions using a single Dynamo file. The spine activity detected from fluorescence recordings was mapped to the reconstructed morphology, allowing synaptic events to be automatically assigned to their corresponding anatomical locations. Detected active spines are highlighted in 3D reconstructions generated from the Dynamo reconstruction using custom visualization scripts (Fig. 7B). The reconstructed morphology was automatically converted into a NEURON model using a custom Python script, and synaptic events identified at active spines were incorporated into the NEURON simulation, with event timings corresponding to the detected calcium peaks from the *in vivo* recordings. Passive and active properties were assigned to dendritic and spine compartments in the NEURON simulation, based on a previously validated model of L2/3 pyramidal cells in VI^35^. Representative simulations generated from the experimentally measured morphology and spine activity are shown in (Fig. 7E-F). These simulations illustrate how dynamic neuronal reconstructions stored in Dynamo can serve as a common interface linking longitudinal structural changes, functional imaging data, and downstream computational modeling within a unified Python workflow. Although demonstrated here using NEURON, the same workflow can be readily extended to other Python-based analysis, simulation, or machine learning frameworks.

**Figure 7.**
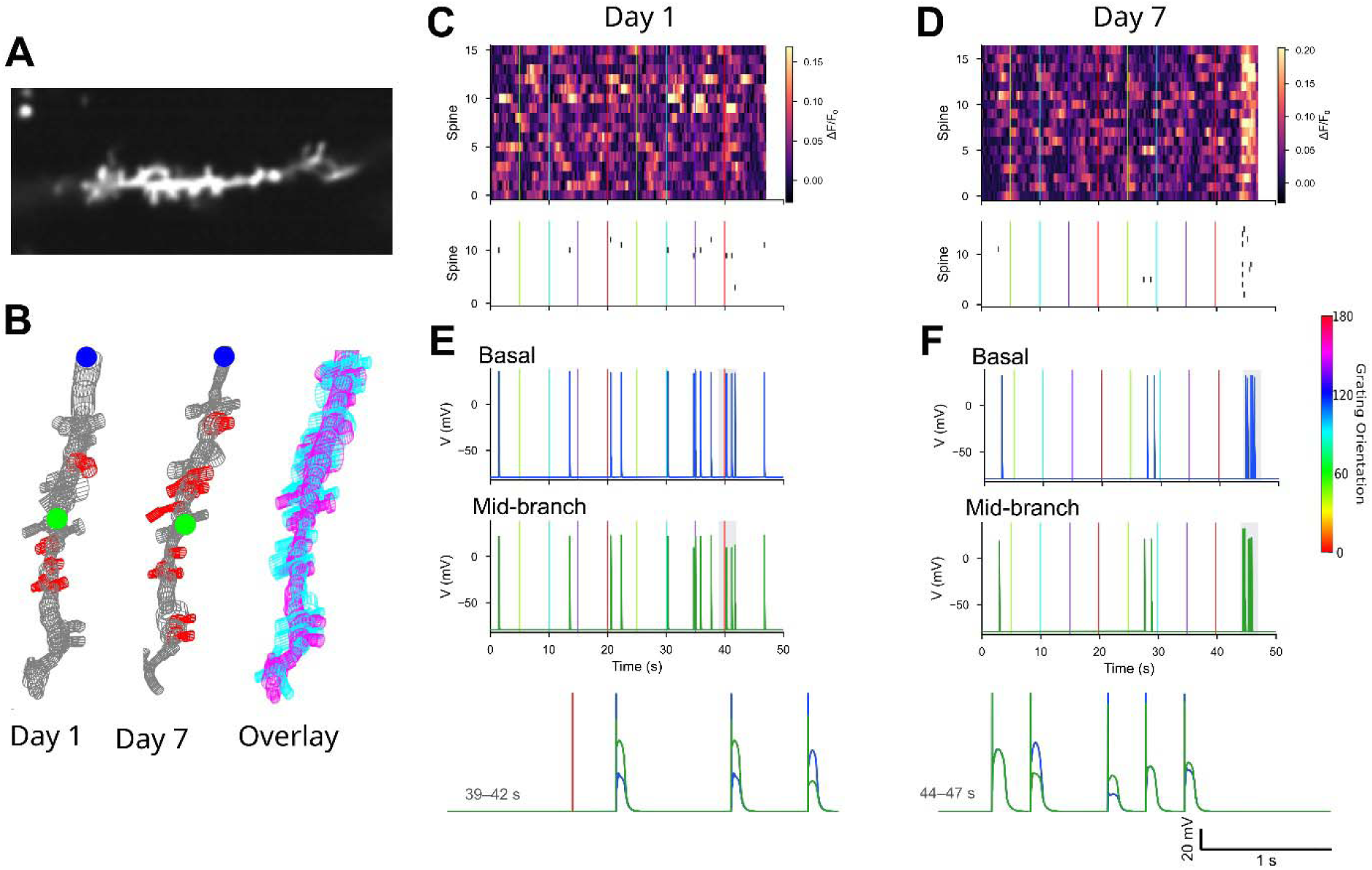
Integration of Dynamo reconstructions linking calcium activity to NEURON simulations. **A,** Dendritic segment on Day 1 (same neuron as in Fig. 6). B, Dynamo 3D reconstructions of the segment in (A) and its spines on Days 1 and 7. Spines with detected calcium events are shown in *red;* the *blue* dot marks the basal recording site used in the simulation and the *green* dot the mid-branch site. At *right,* overlay of the Day 1 *(cyan)* and Day 7 *(magenta)* reconstructions, showing structural change across the imaging interval. **C,** jRGECO1a calcium signal from each spine on Day 1 during visual grating presentation: AF/F_0_ *(top)* and detected events *(bottom).* Vertical lines mark grating onsets, colored by stimulus orientation (color bar). **D,** As in **(C),** for Day 7. **E, F,** Simulated membrane potential at the basal *(blue)* and mid-branch *(green)* recording sites on Day 1, driven by the spine events detected in **(C,D).** Below, the shaded window (39-42 s for Day 1; 44-47 s for Day 7) on an expanded time base with both recording sites overlaid. Color bar indicates grating orientation (0-180°).

## DISCUSSION

Recent advances in sparse-cell labeling and *in vivo* imaging are producing a vast wealth of high-resolution 4D time-lapse images of brain neuronal growth and plasticity creating a need for software capable of efficiently analyzing and quantifying these complex, multi-dimensional datasets. Here, we demonstrate the capabilities of Dynamo, software designed for straightforward reconstruction and rich characterization of neuronal growth dynamics. Dynamo unlocks the full potential of dynamic morphometries by extracting information from digitized neurons in a Python ecosystem, automatically registering the arbor structure and tracking all arbor changes across time. Dynamo reconstructs complete arbors as a hierarchy and automatically synchronizes the identity of every element across a full time series, creating an integrated 4D object that holds identity, topology, and dynamics together. Importantly, given the wide diversity of subject matter and experimental questions which can be addressed using dynamic morphometries, as an open-source platform, Dynamo facilitates versatile modification of analytics to address individual user’s needs allowing for bespoke data pipelines to be readily created and applied.

The need for dynamic morphometric analysis of *in vivo* neuronal time-lapse images is clear from findings that were either challenging or impossible to confirm using static imaging approaches^5^. Examples include: the findings that axonal and dendritic branches can originate from both growth cone splitting and transitions from interstitial filopodia^2,36^; the maturational relationship between dendritic filopodia and spines of different morphologies^37^; identification of new filopodial clustering near previously active synapses^33^; and the establishment of dendritic arbor tiling and polarization^38^. Combining structural imaging with additional markers, such as fluorescently-tagged intracellular proteins, allows for the interrogation of molecular mechanisms underlying dynamic growth and neuronal structural patterning. For example, dual time-lapse imaging of fluorescently-tagged synaptic protein markers along with space-fillers for morphology has provided strong support for the synaptotropic model of axonogenesis and dendritogenesis^39,41^. Further, dual time-lapse imaging of dendritogenesis and calcium biosensors of neuronal activity reveal activity-dependent mechanisms directing structural growth^42^, and the development of faster *in vivo* 4D imaging technologies now opens the possibilities to directly link neuronal information processing and experience-driven functional plasticity to structural plasticity^21^. Dynamo provides an accessible platform for deep analytics of complex structural and functional changes over time.

## Supporting information

4D Dynamo reconstruction tracking dendritic arbor growth

Dual neural structural and activity tracking in Dynamo

Starting a new Dynamo project

Dynamo Drawing Modes

Importing Reconstructions

Arbor Visualization

3D Dynamo reconstruction

## RESOURCE AVAILABILITY

Lead contact: Kurt Haas, PhD;

This study did not generate new materials.

Data and code availability:

Dynamo (RRID: SCR_017541) can be obtained on GitHub: https://github.com/ubcbraincircuits/pyDynamo, and documentation is available at https://ubcbraincircuits.github.io/pyDynamo/.

## ACKNOWLEDGMENTS

This work is supported by funding from a Canadian Institutes of Health Research (CIHR) Foundation Award (FDN-148468) (K.H.), and the Canadian Open Neuroscience Platform (P.C.).

## AUTHOR CONTRIBUTIONS

Conceptualization, P.W.H., P.C., and K.H.; methodology, P.W.H., and P.C.; data analysis, P.W.H., T.D.T., and P.C.; investigation, P.W.H., J.F., and K.P.; writing - original draft, P.W.H., P.C., and K.H.; writing - review & editing, P.W.H. and K.H.; funding acquisition, P.C. and K.H.; supervision, K.H.

## DECLARATION OF INTERESTS

The authors declare no competing interests.

## SUPPLEMENTARY INFORMATION

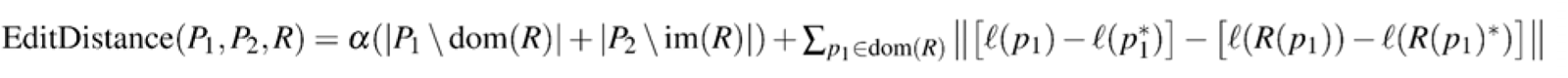

**Sup. Equation 1. Cost Equation for Point ID Registration**. The Registration between two trees is represented by R, an injective partial function registering a set of *Point Objects* between one time point (P_1_) and another (P_2_), potentially leaving some points unregistered. The unmatched error is a, a user-defined parameter controlling how far a point can move before being considered new multiplied by the number of unmatched *Point Objects* in the reference tree (|P1\dom(R)|) and the number of unmatched *Point Objects* in the registered tree (|P2\im(R)|). The distance error calculates how much the point’s location relative to its parent (*l*(p1) - *l*(p1*)) differs between the initial and registered tree (*l*(R(p1)) - *l*(R(p1)*)), and is summed across all registered points.

**Sup. Figure 1.**
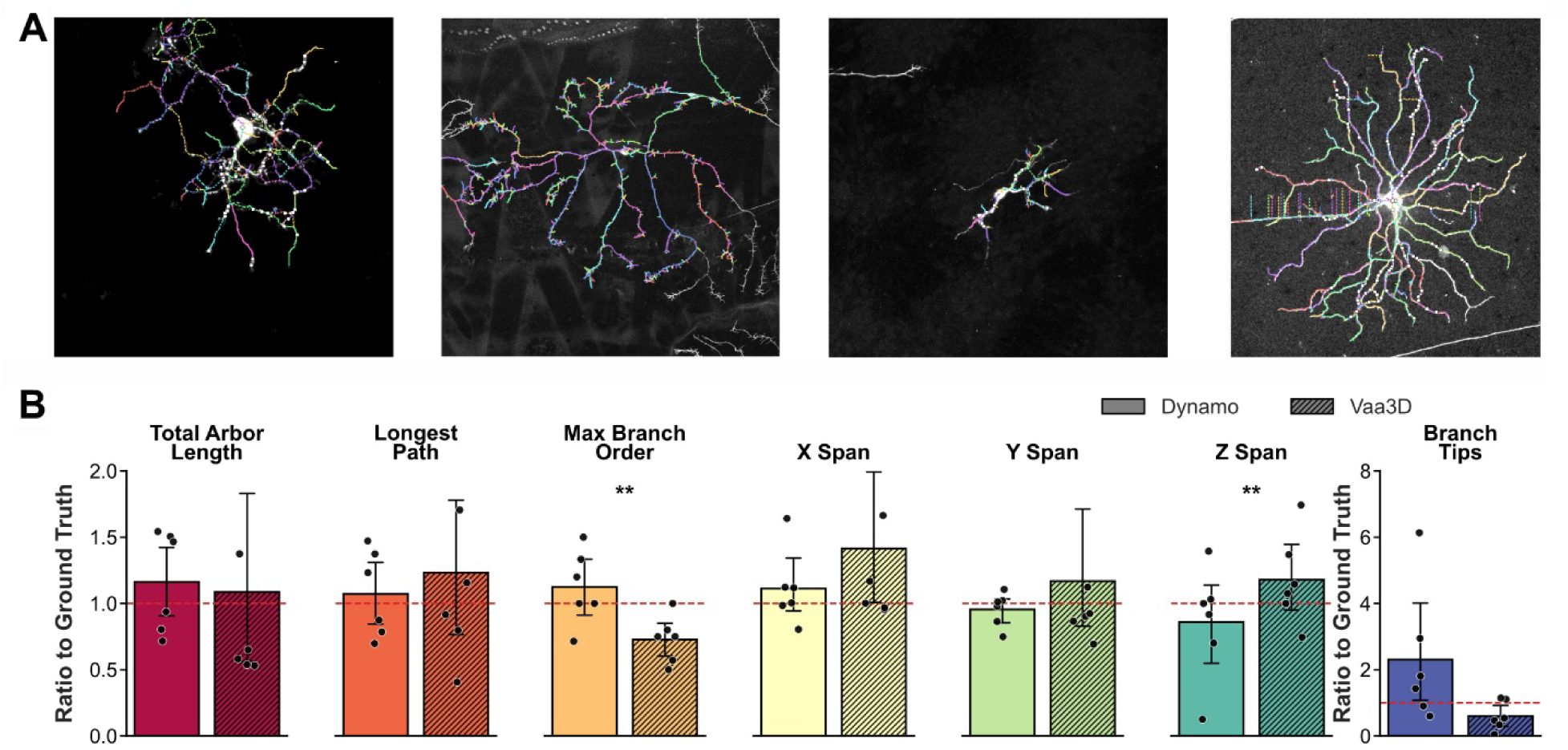
Comparison to modern tracing tools. **A**, Example overlays of Dynamo reconstructions for neurons from *Xenopus, Drosophila,* hIPSCs, and mouse RGCs (left to right), images retrieved from^43^. B, Morphological comparison between Vaa3D and Dynamo traces for neurons from the Goldl66 dataset (N = 6 neurons)^43^ (CI=95%). Statistical tests: paired t-test or Wilcoxon signed-rank test. *p < 0.05, **p < 0.01, ***p < 0.001.

**Sup. Figure 2.**
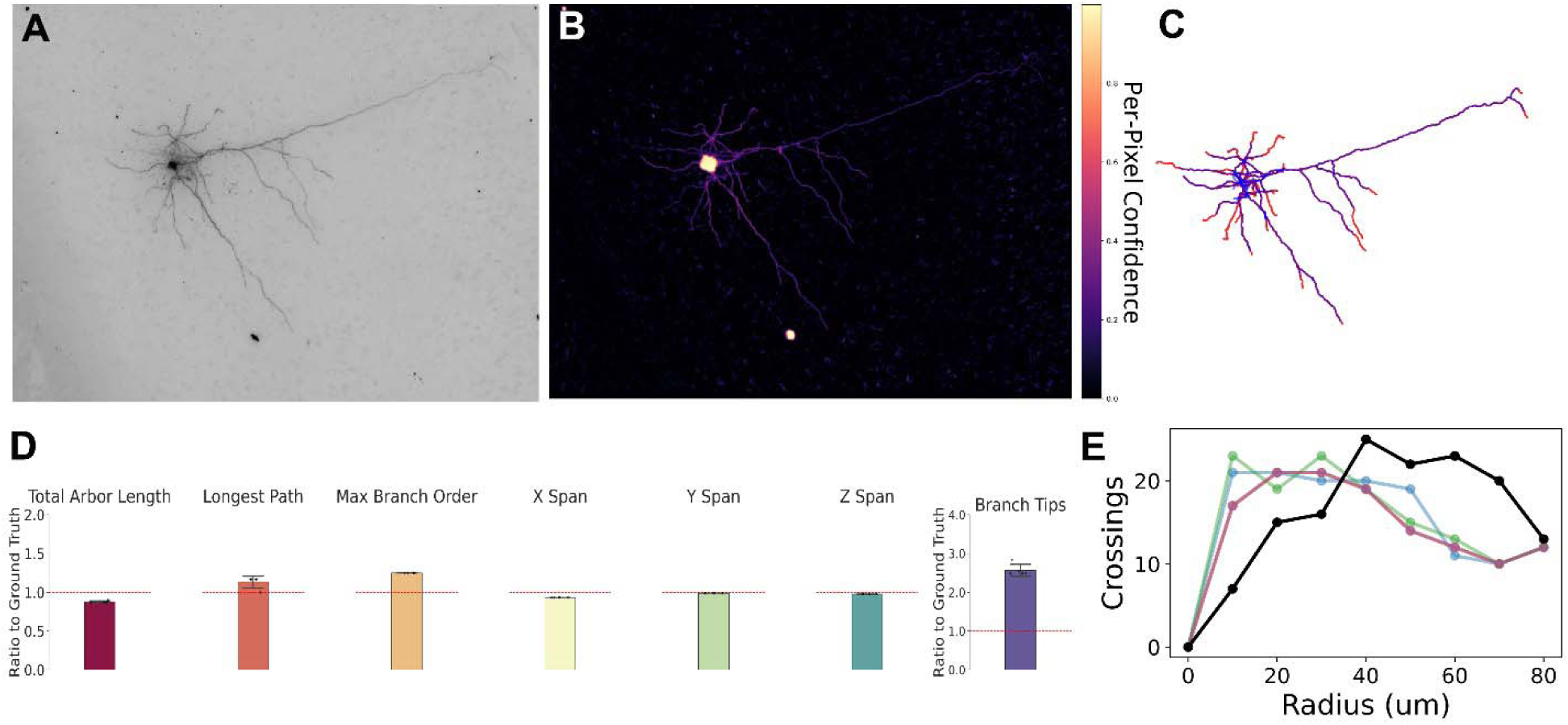
Biocytin tracing from root node. **A**, A minimum intensity projection of the reference image stack. **B**, Pixel-wise confidence map generated by the tracing algorithm, where warmer colors indicate higher confidence. **C,** Reconstruction of the neuronal morphology, with high-confidence regions in purple and low-confidence regions in red. **D**, Morphological features from five independent runs (total arbor length, longest path, maximum branch order, spatial spans in X, Y, and Z, and branch tips) normalized to ground truth values. E, Sholl analysis of the reconstructed neuron showing the number of branch crossings as a function of radius from the soma. Error bars in (D), represent 95% CI (5 runs, N=1 neuron).

**Sup. Figure 3.**
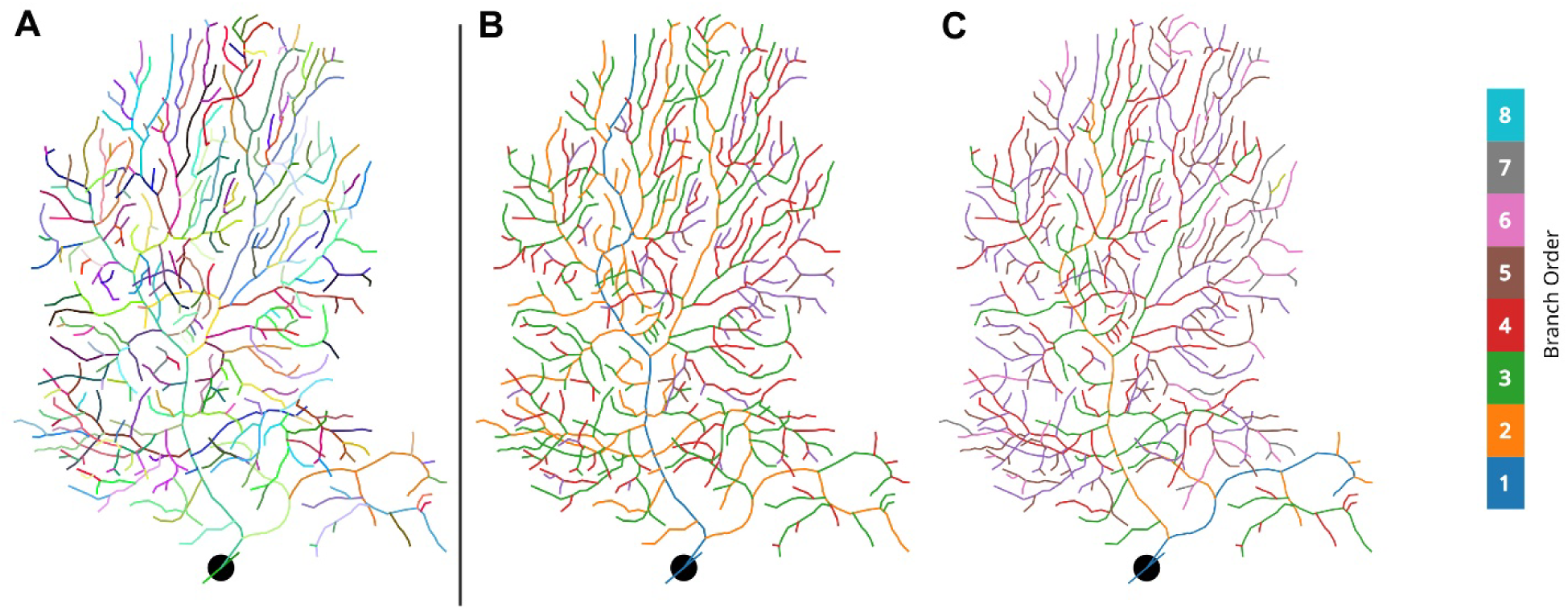
Importing SWC into Dynamo. Dynamo supports importing SWC files to populate the Tree Object with structural information. **A,** Each branch is plotted with a pseudo-random color. **B,C,** Branches can be reparented based either by longest path length (B) or minimum angle (C), with branch color depicting branch order. SWC retrieved from Neuromorpho.org^19^’^44^.

**Sup. Figure 4.**
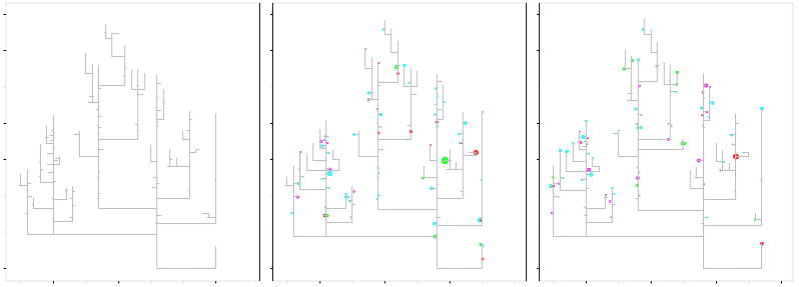
Dendrograms. Dynamo supports displaying growth behavior on flattened dendrograms. These motility diagrams show stable dendritic arbor skeleton in grey and growth behavior across two time points using colored circles at the tips of protrusions displaying new additions *(green),* eliminations *(red),* elongations *(cyan)* and reductions *(magenta),* with circle size depicting change in process length. Left diagram shows dendritic structure at time = 0, middle and right diagrams show growth behavior across two subsequent 30-min intervals.

**Video S1 | Starting a new Dynamo project.** Importing the first image stack of a timelapse using Dynamo’s graphical user interface. The stack is loaded into a Dynamo window where the user can begin manual reconstruction.

**Video S2 | Dynamo Drawing Modes.** The Dynamo GUI supports three different drawing modes. *Skeleton mode* for constructing a wireframe of a neuronal arbor. *Radii mode* to add thickness to the wireframe. And *Puncta mode* to track the location of structure outside of the tree hierarchy.

**Video S3 | Importing Reconstructions.** Reconstructions in the SWC file format can be imported using the Dynamo GUI. Dynamo *Tree Objects* from a proceeding time point can be imported to a newly imported image. After importing the *Tree Object,* image registration can be performed to correct for drift and growth.

**Video S4 | Arbor Visualization.** The Dynamo GUI has several ways of inspecting and visualizing neuronal reconstructions. These include wireframe renderings and volumetric renderings. Both can be used to inspect the quality of a reconstruction

**Video S5 | 3D Dynamo reconstruction.** A single time point Dynamo reconstruction of a Xenopus tectal neuron imaged in vivo and rendering in Napari. The movie transitions from the raw 3D image volume encompassing the entire soma and dendritic arbor, followed by an overlay of a Dynamo-generated 3D digital reconstruction, and then by the reconstruction alone.

**Video S6 | 4D Dynamo reconstruction tracking dendritic arbor growth.** Time series of Dynamo digital reconstructions of a *Xenopus* tectal neuron next to the reference stacks.imaged at 3-min intervals, with growth behavior of dendritic filopodia (branches < 10pm) depicted by colors: new additions *(green),* eliminations *(red),* elongations *(cyan)* and reductions *(magenta).* Movie rendered in Napari^29^.

**Video S7 | Dual neural structural and activity tracking in Dynamo.** *In vivo* imaging of visual-evoked neuronal activity at each spine in an optical plane of a mouse VI pyramidal cell are tracked between two imaging sessions spaced a week apart. *Top panels:* Wireframes of Dynamo reconstructions are plotted over the reference images, with the color of each spine based on the *Branch Object* ID. A circle is placed at the tip of each spine, with circle size and color corresponding to dF/F value of the jRGECO1 calcium sensor signal from ROI generated using the Dynamo reconstruction. *Bottom panels:* Calcium sensor traces for the population average of all spines sampled *(grey trace)* and an individual spine indicated by a yellow arrow *(red trace)* are displayed for each day.

**Sup. Table 1.**
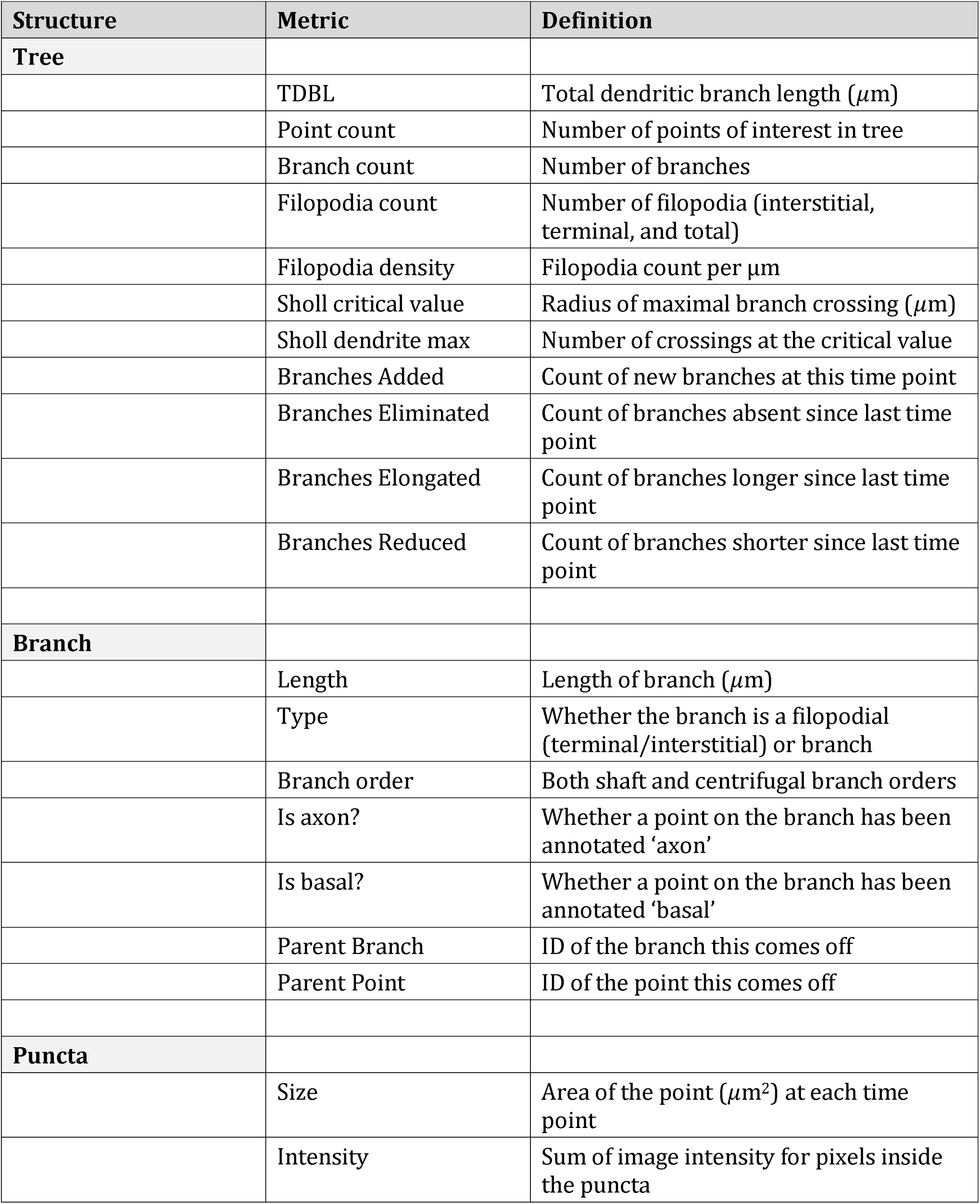
Supported Export Metrics

## STAR METHODS

### EXPERIMENTAL MODEL and STUDY PARTICIPANT DETAILS

#### Animals

The imaging volumes in this paper were recorded from a stage 48 albino *Xenopus laevis* tadpole, reared at 22°C in 10% Steinberg’s solution on a 12-hr light/dark cycle. All experimental procedures were conducted according to the guidelines of the Canadian Council on Animal Care, and were approved by the Animal Care Committee of the University of British Columbia.

### METHODS DETAILS

#### *In Vivo* Single Cell Labeling

The *Xenopus laevis* optic tectum neuron imaged was transfected *in vivo* with farnesylated enhanced green fluorescent protein (EGFP) by single-cell electroporation^8^. Electroporation was conducted under anesthetic with 0.02% MS-222 using plasmid DNA (1 Mg/MD and an Axoporator 800A stimulator (Molecular Devices, Sunnyvale, CA) to deliver electric pulses (stimulus parameters: pulse intensity = 1.5 *pA;* pulse duration = 1 ms; pulse frequency = 300 Hz; train duration = 300 ms).

#### *In Vivo* Time-lapse Imaging of Dendritogenesis

Image volumes of the *Xenopus* tectal neuron were acquired using *in vivo* time-lapse microscopy with a custom-built two-photon laser-scanning microscope (TPLSM) made by sending a Chameleon XR laser (Coherent, Santa Clara, CA) through an Olympus FV300 confocal microscope (Olympus, Center Valley, PA), and image captured using a 60X, 1.1 numerical aperture, water-immersion objective (LUMPlanFI, Olympus), and Fluoview software (Olympus). The tadpole was paralyzed by a 5 min exposure to 3 mM pancuronium dibromide (PCD), and continuously perfused with oxygenated Steinberg’s solution 22°C. A volume containing the arbor was repeatedly captured at 3-min intervals over 30 min (LifeActEGFP) or 5-min intervals over 120 min (EGFP) using a z axis step of 1.5 μm, a resolution of 512 x 512 px and zoom factor of 1.5X, encompassing a region of interest of 157 x 157μm.

#### Machine Learning-Based Automated Neural Reconstruction

We trained a U-Net with residual blocks on individual slices of 3D image stacks to classify pixels as either soma, dendritic arbor, or background autofluorescence^47,46^. The training dataset consisted of 20 EGFP-expressing tectal neurons imaged as described above, and with pixel classification annotated by experts to generate a ground truth dataset, resulting in a training set of 1294 2D slices. Raw image pixel intensity values were normalized between 0 and 1 for numerical stability during training. To enhance training and reduce overfitting, the training set was increased by augmenting image slices containing labels using the TorchIO package, increasing the overall number of training images^47^. This involved adjusting the training images with gamma values ranging from (-0.6,0.5), injecting gaussian noise with a mean of 0.05 and a standard deviation of 0.025, as well as rotating the slices about the axial dimension between (-25,25) degrees. Ground truth images were split into 75% training and 25% validation sets.

The network was trained with the following hyperparameters: 2 residual units, 5 downsampling and upsampling layers with number of filters starting from 32 to 512 filters, doubling the filters every subsequent layer using 3x3 filter size with stride 2, P-ReLU for activation layer, batch normalization, and a dropout rate of 0.10 to prevent overfitting^45^. The training process utilized a combination of dice and categorical cross entropy loss for 80 iterations with the AdamW optimizer with a weight decay of 0.00421^48,49^. We also used a cosine annealing learning rate scheduler with an initial learning rate of 7.54e-5, resetting the learning rate every 20 epochs. Hyperparameters such as the learning rate, dropout, and weight decay were tuned using Bayesian optimization^50^. The Residual U-Net was implemented using the MONAI package^51^, and the training was performed on Compute Canada high-performance computing cluster, Cedar, using an NVIDIA V100 Large GPU. Pixels classified as arbor were skeletonized in 3D for the entire volume. The binary skeleton was segmented into individual branches using a kernel to identify branching nodes and branch ends. Points belonging to each branch were ordered by distance from either a branch end or branching node. The tree structure was formed using a nearest-neighbor algorithm to first connect the branches of basal and primary dendrites, identified by their proximity to the soma. Additional branches were attached using a nearest neighbor algorithm between the first point of the branch segment and the pool of points already in the tree. Once the tree was assembled the point density of individual branches was reduced with a 3D implementation of the Ramer-Douglas-Peucker algorithm, smoothing the tree structure. This method of neuronal reconstruction was compared to the ground truth reconstructions from a published dataset^43^. Inclusion criteria that the U-Net could detect the neuron’s soma and that the APP2 algorithm from Vaa3D could trace the neuron with its default parameters were used.

A second automated reconstruction method was developed for mammalian cells labeled with biocytin. Images of neurons labeled with biocytin were retrieved from the Allen Institute’s Allen Cell Types Database using an API^52^. Image stacks consisted of hundreds of slices and were several thousand pixels in the X and Y dimensions. To reduce computational load during processing, images were down sampled laterally by a factor of 4 using the mean pixel intensity from non-overlapping 4x4 blocks.

Four neurons from the database were used as ground truth to train a 2D segmentation model. The ground truth was generated using the SWC file for the corresponding neuronal image stack. SWC coordinates were scaled into pixel space to match their corresponding reference images. For each line segment in the SWC file, the corresponding labeled pixels in a label array were filled using Bresenham’s line algorithm^53^ and dilated using the radius value in the SWC file to match the radius of the neuron arbor. The labels were further dilated and collapsed from the four adjacent z-planes to ensure complete coverage of the neuronal pixels in the reference images, as branches frequently had signal in multiple image planes. These automated labels were then manually cleaned using Napari to remove errors and improve the quality of the training data. Pairs of 64x64 images and labels were randomly sampled around the neuronal structure to generate a training set; these images were only kept if they contained neuronal labels. This resulted in a total of 48,738 images in the training set, of which 80% were used for training and the remainder for validation.

The neuronal arbor was assembled using the same method as above, with one difference, the root-node is provided by the user.

#### Imaging and Analysis of Dendritic Spine Plasticity in Mouse Visual Cortex

The mouse visual cortex data re-analyzed here were originally acquired by K.P. at the HHMI Janelia Research Campus and are included in the supplementary material of ref 21. Briefly, neurons in the left visual cortex of a C57Bl/6NCrl mouse were sparsely labeled with two AAVs encoding jRGECO1a and iGluSnFR-SF-VenusA184S and a third encoding Cre as previously described^21^. All animal procedures were in accordance with protocols approved by the HHMI Janelia Research Campus Institutional Animal Care and Use Committee (IACUC 17-155). Two imaging sessions were conducted seven days apart while mice were lightly anaesthetized with vaporized isoflurane (0.75% V/V) and presented moving-bar visual stimuli as previously described^21^. A 3D image volume was collected, followed by 2D functional imaging of visual responses to 8 directions of motion stimuli (8 repetitions each). Spines were classified using established categories that were defined using measurements of spine width and length taken directly from Dynamo reconstructions^29^. Activity traces from spines were generated using a script that converted Dynamo reconstructions into regions of interest to measure fluorescent intensities from individual spines in two color channels of an *in vivo* recording. Traces were saved in NWB format and imported into the Dynamo project file^31^.

#### Biophysical Simulation

Dynamo structures were parsed into a NEURON (8.2.6+) using a custom Python script, which used one branch from the reconstructed neuron from Figure 7 to generate the structural data for the simulation. Using the calcium traces generated from individual spines, synaptic events were detected using a threshold method. The 5F/F was calculated using methods implemented in PyNeurotrace^32^. Events were defined as peaks in the calcium traces with an amplitude four standard deviations or higher of calcium trace.

Spines with detected events were given synapses with both AMPAR (rising time constant of 0.1 ms, and decay of 2.5 ms) and NMDAR (rising time constant of 2 ms, and decay of 30 ms) synaptic inputs. The active properties of the dendrite used a previously published model of layer2/3 pyramidal cells from VI^35^. The corresponding files for the model were retrieved from ModelDB, model number 267501^54^. The simulation was run for 60000 ms with a time step of 0.1 ms. The membrane potential of the dendritic branch was recorded from the first segment near the base of the branch, and the middle segment of the main branch.

#### Quantification and Statical Analysis

The sample size of independent biological replicates, neurons measured, (N) is provided in each figure legend. A paired t-test or Wilcoxon signed-rank test were used with *p <0.05, **p < 0.01, ***p < 0.001. Results are presented as mean ± SEM.

#### Key Resources Table

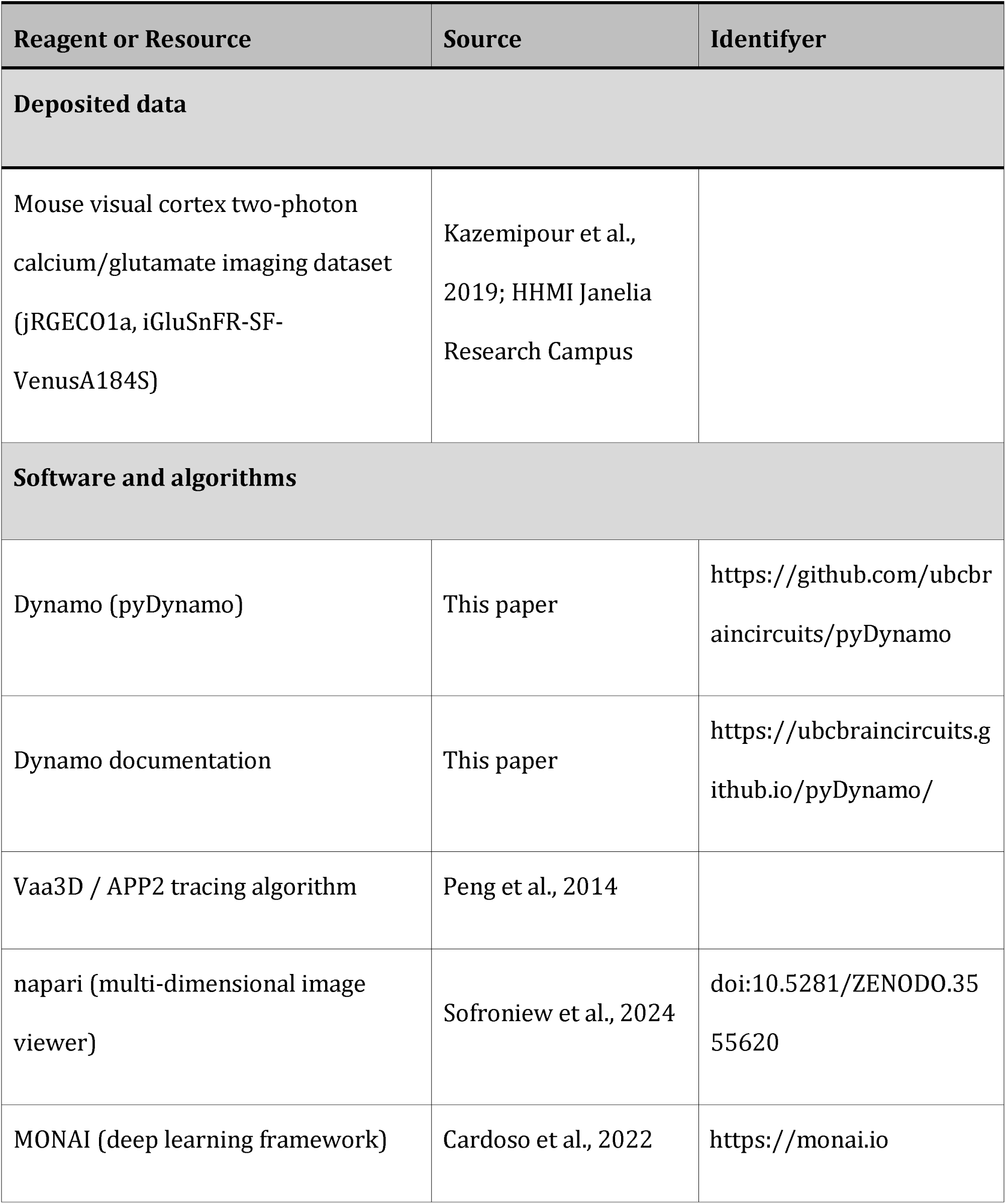
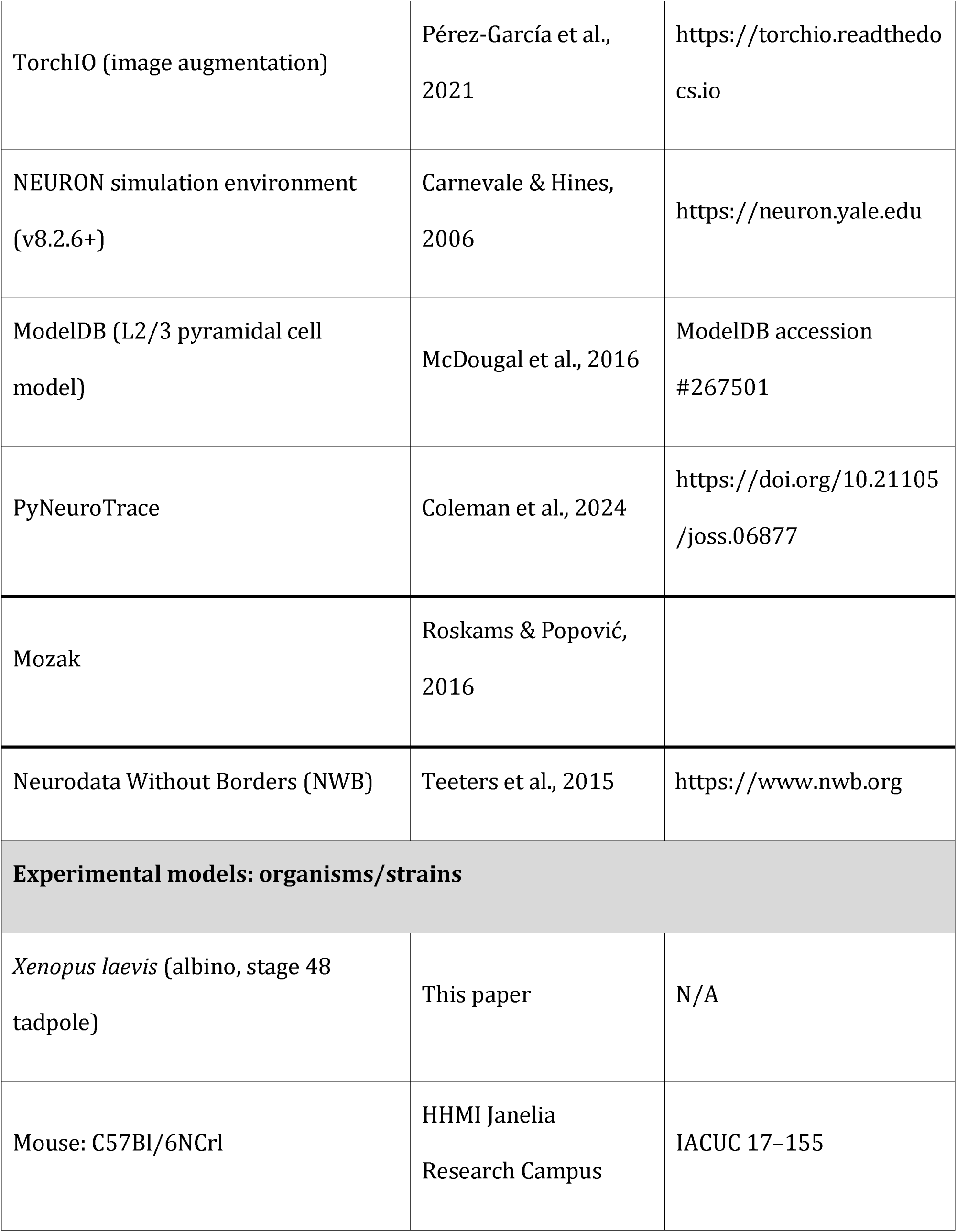
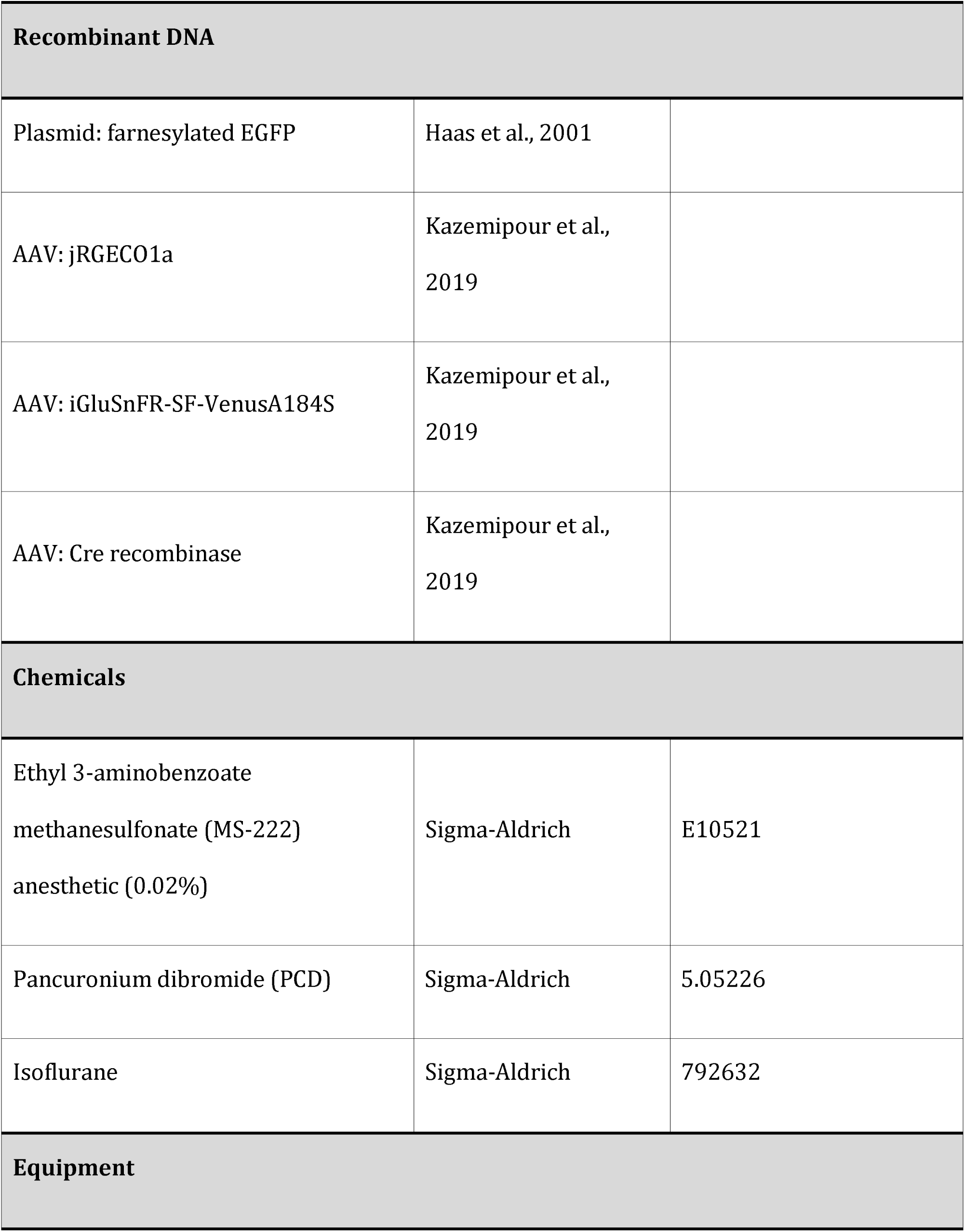
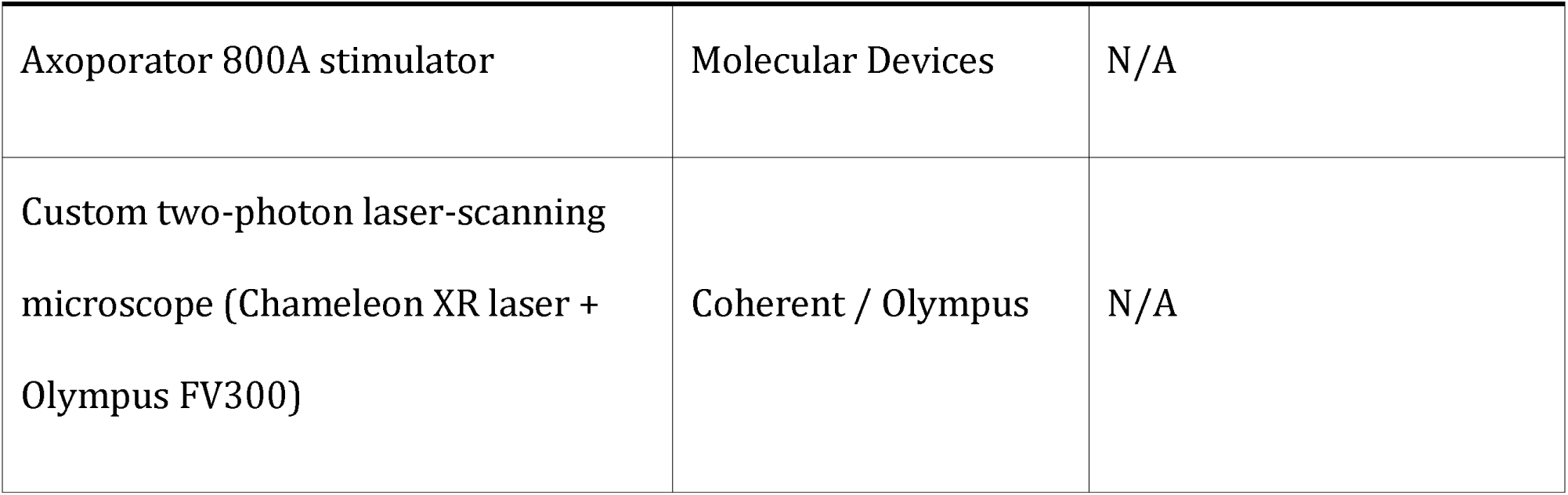

## Notes

https://github.com/ubcbraincircuits/pyDynamo

### Competing Interest Statement

The authors have declared no competing interest.

